# A consortium-based approach to adaptive laboratory evolution of *Acinetobacter baylyi* ADP1 for lignin valorization

**DOI:** 10.1101/2024.08.26.609767

**Authors:** Suchismita Maiti, Prashant Singh, J Vishnu Prasad, Anantha Barathi Muthukrishnan, Lars M. Blank, Guhan Jayaraman

**Affiliations:** Department of Biotechnology, Bhupat and Jyoti Mehta School of Biosciences, Indian Institute of Technology Madras, Chennai, Tamil Nadu, 600036, India

## Abstract

The utility of *Acinetobacter baylyi* ADP1 (ADP1) for lignin valorization has yet to be sufficiently investigated compared to other organisms such as *Pseudomonas*, *Rhodococcus,* etc. In this study, a two-step Adaptive Laboratory Evolution (ALE) process was used to evolve a unique ADP1 strain (*A. baylyi* SAG_185). Initially, several ADP1 strains were evolved for substrate tolerance to specific lignin-related aromatics (LRAs). Subsequently, a consortium of these strains was adaptively evolved in a mixture of LRAs, which resulted in the evolution of SAG_185. This strain was capable of simultaneous utilization of multiple LRAs at higher concentrations as well as grow on a depolymerized lignin-rich residue obtained from enzymatic hydrolysis of pre-treated corncob. This is the first report on such an evolutionary strategy.

Whole-genome sequence analysis of all the evolved strains revealed large-scale mutations involving insertion sequences (IS). In particular, SAG_185 revealed a critical mutation in the vanR repressor gene, resulting in the up-regulation of vanAB genes required to convert vanillate to the key intermediate, protocatechuate (PCA). Additionally, there were two large deletions of 9kb and 38kb DNA segments, including genes for putative transcriptional regulators of LysR, MarR and AraC family. The evolved strains also showed mutations in the hcaE gene, responsible for the uptake of LRAs. The vast number of mutations in hypothetical proteins, transporter and regulatory sequences indicate the underlying effects of these regions on the uptake of multiple LRAs. Overall, our findings provide potential targets for reverse engineering of A. baylyi ADP1 for lignin valorization.

**IMPORTANCE:** This study shows a novel strategy for adaptive laboratory evolution, which can be generically adopted to evolve bacterial strains for taking up multiple substrates which are toxic at higher concentrations. We developed a two-step evolutionary strategy to evolve a unique strain, A. baylyi SAG_185, which could take up multiple lignin-related aromatic monomers at higher concentrations as well as grow on depolymerized lignin. Initially, individual strains were adapted to utilize single aromatic monomers at higher concentrations. After many unsuccessful attempts to adapt these strains for utilizing multiple monomers, a consortium of the five evolved strains were grown on a mixture of aromatics and adapted to utilize all the monomers at high concentrations. The adapted consortia resulted in the evolution of SAG_185. Whole-genome sequence analysis of all these strains gave rise to many interesting insights on potential genetic targets for reverse engineering of A. baylyi ADP1 for lignin valorization.

## INTRODUCTION

*Acinetobacter baylyi* ADP1 (ADP1) is a strictly aerobic Gram-negative soil bacterium capable of degrading aromatic compounds through the β-ketoadipate pathway (1). ADP1 has drawn considerable attention as a desirable host for metabolic engineering due to its compact genome size (3.6 Mb), natural competence, high rate of homologous recombination, and the availability of genetic tools for genome engineering (2–4). Many researchers have engineered ADP1 to produce compounds such as cyanophycin (5), alkane (6), wax esters (6, 7), triacylglycerols (8), 1-alkene (9), poly-α-olefins (10), mevalonate (11), and naringenin (12). *Acinetobacter* species are also known to degrade various aromatic compounds like chlorobenzoate (13), xenobiotics such as benzoates, toluene, phenol (14) and lignin-related aromatics (LRAs) such as trans-ferulic acid and p-coumaric acid (15).

Lignin is a recalcitrant component of lignocellulosic biomass. Depolymerization of lignin results in many aromatic monomers which are difficult to valorize using microbial strains. These aromatics can be converted to many value-added compounds such as cis, cis-muconic acid (16) and protocatechuate (17). Most of the researchers have used thermo-chemical depolymerization methods (18–21) to produce lignin-derived monomers (LDMs) such as p-coumaric acid, ferulic acid, vanillic acid, and 4-hydroxybenzoic acid. Phenolic compounds from lignocellulosic hydrolysates are toxic to microorganisms (22–24). However, wild-type ADP1 can tolerate LRAs (6, 9, 11), although it is sensitive to these LRAs beyond certain concentrations. Microbial strains need to adapt to such toxic compounds at high concentrations for their effective utilization from depolymerized lignin.

To overcome the above challenges, strain improvement is essential for enhanced tolerance to LRAs and inhibitors like furfural and 5-(hydroxy-methyl)-2-furaldehyde (25) ALE has been used to adapt ADP1 on different carbon sources, such as terephthalic acid (26) and ferulate (9, 27). Similarly, ALE improved the tolerance and catabolism of *Pseudomonas putida* KT2440 towards ferulic acid and *p*-coumaric acid (28).

ALE has often resulted in identification of non-obvious, novel targets for reverse metabolic engineering. Previous ALE studies with ADP1 have focused on the uptake of individual LRAs such as ferulate (9, 27). Reverse metabolic engineering of ADP1 was enabled by whole-genome sequence analysis of ferulate-evolved strains which revealed critical mutations in the aromatics-transporter genes (*hcaE*, *hcaK* and *vanK*). It is also important to de-bottleneck the metabolic pathways for utilization of aromatics such as ferulate, p-coumarate, vanillate, caffeate and 4-hydroxybenzoate which are processed through the PCA branch of the β-ketoadipate pathway in ADP1.

In this study, we developed a two-stage adaptive evolution strategy to evolve a unique strain capable of metabolizing multiple LRAs as well as grow on depolymerized lignin. Initially, we subjected ADP1 to ALE on a single LRA and evolved five different strains adapted to five synthetic LRAs, i.e., p-coumaric acid, ferulic acid, vanillic acid, caffeic acid and 4-hydroxybenzoic acid, with each strain having tolerance to an elevated concentration of the respective LRA (Table 2). Since the evolved ADP1 strains were unable to utilize multiple LRAs at high concentrations, we grew a consortium of the five evolved ADP1 strains on a mixture of the LRAs. The consortium was adapted to an increasing concentration of all the LRAs in this mixture. This approach resulted in the evolution of a single unique strain which was able to efficiently utilize all the five LRAs simultaneously. To understand the genetic basis of these adaptations, whole-genome sequencing (WGS) and comparative analysis was performed for these strains to identify gene targets for reverse metabolic engineering of ADP1.

## MATERIALS AND METHODS

### Bacterial strain and growth conditions

Wild-type ADP1 strain was obtained from DSMZ, Germany (DSM No. 24193). Minimal Salt Medium (MSM) (6) was used for all the experiments containing (L^-1^); K_2_HPO_4_ -3.88 g, NaH_2_PO_4_ -1.63 g, (NH_4_)_2_SO_4_ - 2.00 g, MgCl_2_·6H_2_0 - 0.1 g, EDTA - 10 mg, ZnSO_4_·7H_2_O - 2 mg, CaCl_2_·2H_2_O - 1 mg, FeSO_4_·7H_2_O - 5 mg, Na_2_MoO_4_·2H_2_O - 0.2 mg, CuSO_4_·5H_2_O - 0.2 mg, CoCl_2_·6H_2_O - 0.4 mg, MnCl_2_·2H_2_O - 1 mg. A stock solution of 2 M glucose was used as a carbon source whenever required. Five different synthetic LRAs, viz. p-coumaric acid (Cm), 4-hydroxybenzoic acid (Hb), vanillic acid (Vn), ferulic acid (Fe) and caffeic acid (Cf) (Sigma Aldrich) were used for all the ALE studies. Initially 0.5 M stocks of all the 5 LRAs were prepared by dissolving them in 2 M NaOH. Three different synthetic-LRA mixtures were prepared for ALE on mixtures, as mentioned in Table 1. All growth conditions were maintained at 275 rpm, pH 7 and 30 °C.

**Table 1.** Media compositions containing mixture of LRAs.

| Media name | Concentration of each LRA in the mixture (mM) |  |  |  |  |
| --- | --- | --- | --- | --- | --- |
|  | p-Coumaric acid | 4-Hydroxybenzoic acid | Caffeic acid | Ferulic acid | Vanillic acid |
| MM-I | 20 | 20 | 5 | 5 | 10 |
| MM-II | 25 | 25 | 10 | 10 | 15 |
| MM-III | 25 | 25 | 15 | 15 | 20 |

### Adaptive laboratory evolution (ALE) on individual LRAs

Initially, the ADP1 strain was screened for the tolerance levels of each individual LRA by plating in minimal media. The ALE process was based on the technique described by Kurosawa et al. (2015) (22). The initial concentrations of the LRAs (Table 2) were selected based on previous literature (6, 9, 29–31). A 10% v/v inoculum from the previous culture was sequentially transferred to fresh media (5 mL) and sub-cultured at a particular concentration of the LRA till the highest growth rate was achieved (Supplemental Material 1 Table S2). When no further improvement in growth rate was observed, the culture was transferred to a media containing a higher concentration of LRA. This process was repeated by sequentially increasing the concentration of each LRA by 5mM, till the final concentration (Table 2), when further growth was not possible.

**Table 2.** The minimum and maximum concentrations of the LRAs used to adapt different strains.

| LRAs | Evolved ADP1 Strains | Initial Concentration (mM) | Final Concentration (mM) |
| --- | --- | --- | --- |
| p-Coumaric acid (Cm) | SAG_Cm | 20 | 60 |
| 4-Hydroxybenzoic acid (Hb) | SAG_Hb | 20 | 100 |
| Caffeic acid (Cf) | SAG_Cf | 5 | 45 |
| Ferulic acid (Fe) | SAG_Fe | 5 | 50 |
| Vanillic acid (Vn) | SAG_Vn | 10 | 100 |

### ALE on synthetic LRA mixtures

After adaptation of strains to higher concentrations of the individual LRAs, the evolved strains i.e., SAG_Cm, SAG_Hb, SAG_Vn, SAG_Fe and SAG_Cf were grown on a synthetic mixture of LRAs (MM-I). None of the above strains, which were adapted to a particular LRA, showed the ability to utilize other LRAs in the mixture. Therefore, we attempted to further evolve each of these strain by sequentially adapting them to an additional LRA. Out of the 20 strains we attempted to evolve, 3 showed stable growth and could simultaneously utilize 2 LRAs at higher concentrations. In a subsequent set of experiements, a consortium of the five single-LRA- evolved strains were grown in the media MM-I and the consortium was further evolved in MM-II containing higher concentrations of all the LRAs (Table 1). This experiment was repeated several times, and we found three consortia which were adapted to MM-II and effectively consumed all the LRAs in MM-II. After plating the evolved consortium of strains, we could isolate a single unique strain (SAG_185), whose genotypic and phenotypic characteristics are described in detail in the Results and Discussion sections.

### Base-catalyzed depolymerization of lignin from pre-treated corncob

A lignin-rich fraction (50-55% lignin) was obtained from the cellulosic hydrolysate of pre-treated corncob. This was subjected to base-catalyzed depolymerization (BCD) using 4% NaOH with a 10% w/v (50 mL) biomass in a 250 mL Schott DURAN® bottle. The depolymerization was performed in an autoclave (Equitron®, India) under the operating conditions of 40 min and 121 °C followed by centrifugation (Eppendorf 5810 R) at 8000 g for 20 mins. The supernatant was separated and analyzed using HPLC. For growth studies, the extracted supernatant was added to the media along with other MSM media components. The media was diluted to keep p-coumaric acid to 20 mM and the final pH was adjusted to 7. This media was named as BCD_Cm20.

### Growth and substrate utilization studies

All studies on growth and substrate utilization were performed in 50 ml cultures (250 ml shake flasks). To estimate biomass, the optical density was measured at 600 nm (OD600) using a spectrophotometer (Jasco Corp. V-550). The LRAs and protocatechuate were analysed by HPLC (Shimadzu) on a Luna C18 column (Phenomenex – 250 x 4.6 mm, 5 µm particles), using the protocol described by Schwarz et al. (2009) (32). The chromatographic conditions were: 1 mL/min flow rate; 20 μL injection volume and eluents: A (5% methanol, 2% acetic acid, 93% water) and B (90% methanol, 2% acetic acid, 8% water).

### Whole-genome sequencing and analysis

The ALE processes described above resulted in unknown genetic changes in the evolved strains. Therefore, the genomic DNA of each evolved strain was isolated using the Qiagen QIAamp® DNA Mini Kit according to the manufacturer’s protocol for WGS studies. WGS of seven ADP1 strains (ADP1, SAG_Cm, SAG_Hb, SAG_Vn, SAG_Fe, SAG_Cf, and SAG_185) were performed by Macrogen Inc. (South Korea). The sequencing was done using Illumina sequencing by synthesis (SBS) technology, which involved short read (101-151 bp) paired-end sequencing.

The raw sequence data were analyzed using breseq (version 0.35.6) (33) in consensus mode with default consensus and polymorphism frequency cut-offs of 0.8 and 0.2, respectively. The workflow pipeline utilized R (version 3.6.3) and Bowtie (version 2.3.5.1). The ADP1 complete genome (GenBank: CR543861.1), which is IS-element annotated by Dr Jeffery E. Barrick’s Lab (UT Austin), was used as a reference genome. The sequence alignment was performed using the "bwa-mem" algorithm (34) and the percentage mapping was calculated using the SAMtools-1.10 "flagstat" option tool (35).

### RNA isolation and reverse transcription

Following the manufacturer’s protocol, the RNeasy Plus Mini Kit (QIAGEN) was used to extract total RNA at mid-log phase from all the strains. RNA concentration and purity was assessed using a Denovix DS-11 spectrophotometer. To evaluate the integrity of ribosomal RNA bands, 1.2% agarose gel electrophoresis was performed. Starting with 1 μg of RNA, a reaction for reverse transcription was performed using the RT2 First Strand Kit (QIAGEN).

### qPCR

The forward and reverse primers for all the target genes (*vanA*, *vanB*, and *vanR*) and reference genes (*ileS* and *rpoB*) are listed in the Supplemental Material 1 (Table S6). These primers were designed using the "Benchling" software and the primer efficiencies were calculated using standard curve method described by Svec et al. (2015) (36). All the real-time qPCR studies were performed using 0.2 ml 96-well plates QuantStudioTM 3 Real-Time PCR (Applied Biosystems) using the following temperature profile: 95 °C for 30 s, and 40 cycles of 95 °C for 5 s and 58 °C for 30 s. The reaction volume was 20 μL including 0.8 μL forward primer (10 μM), 0.8 μL reverse primer (1 μM), 0.4 μL ROX Reference Dye (50X), 10 μL TB Green Premix Ex TaqTM II (Tli RNaseH Plus) (2X), 6 μL nuclease-free water (QIAGEN) and 2 μL template cDNA. For no template control (NTC), instead of template cDNA, nuclease-free water was added.

### Statistical analysis of qPCR data

The relative gene expression (RGE) was calculated using the modified Pfaffl (2001) (37) method by taking the geometric mean of the relative quantity (RQ) of two reference genes. ADP1 was used as the calibrator sample. The qPCR experiment was conducted with biological triplicates and two technical replicates to ensure statistical significance. Analysis of variance (ANOVA) was performed at a 95% confidence level to assess the statistical difference in gene expression across different strains. Subsequently, Tukey’s Honestly Significant Difference (HSD) test, was conducted to determine the significant differences in gene expression between the strains.

## RESULTS

### ALE of ADP1 strains on individual LRAs

For conducting ALE studies, five different LRAs were chosen based on the PCA branch of β-ketoadipate pathway (Fig. 1), viz. caffeic acid, p-coumaric acid, 4-hydroxybenzoic acid, ferulic acid and vanillic acid. Initially, the wild-type ADP1 was subjected to a toxicity check on individual LRAs present in minimal media plates. The strains exhibited growth on plates containing individual LRAs, with the toxicity limits for the LRAs ranging from 25-55 mM (Supplemental Material 1 Fig. S1). Subsequently, the first stage of ALE involved successive adaptation of wild-type ADP1 to gradually increasing concentrations of individual LRAs mentioned above. This resulted in the evolution of five different strains of ADP1, each adapted over multiple generations (Supplemental Material 1 Table S1) to a different maximum concentration of the individual LRAs ranging from 45 mM (caffeic acid) to 100 mM (4-hydroxybenzoic acid and vanillic acid) (Table 2) The growth and substrate utilization for the 5 strains are provided in Supplemental Material 1 (Table S2). However, none of these strains could grow in higher concentrations of the other LRAs, i.e. apart from the one on which they were adapted (Supplemental Material 1 Fig. S2). This indicates that the evolution of each of these strains were specific to only one LRA.

**Fig. 1.**
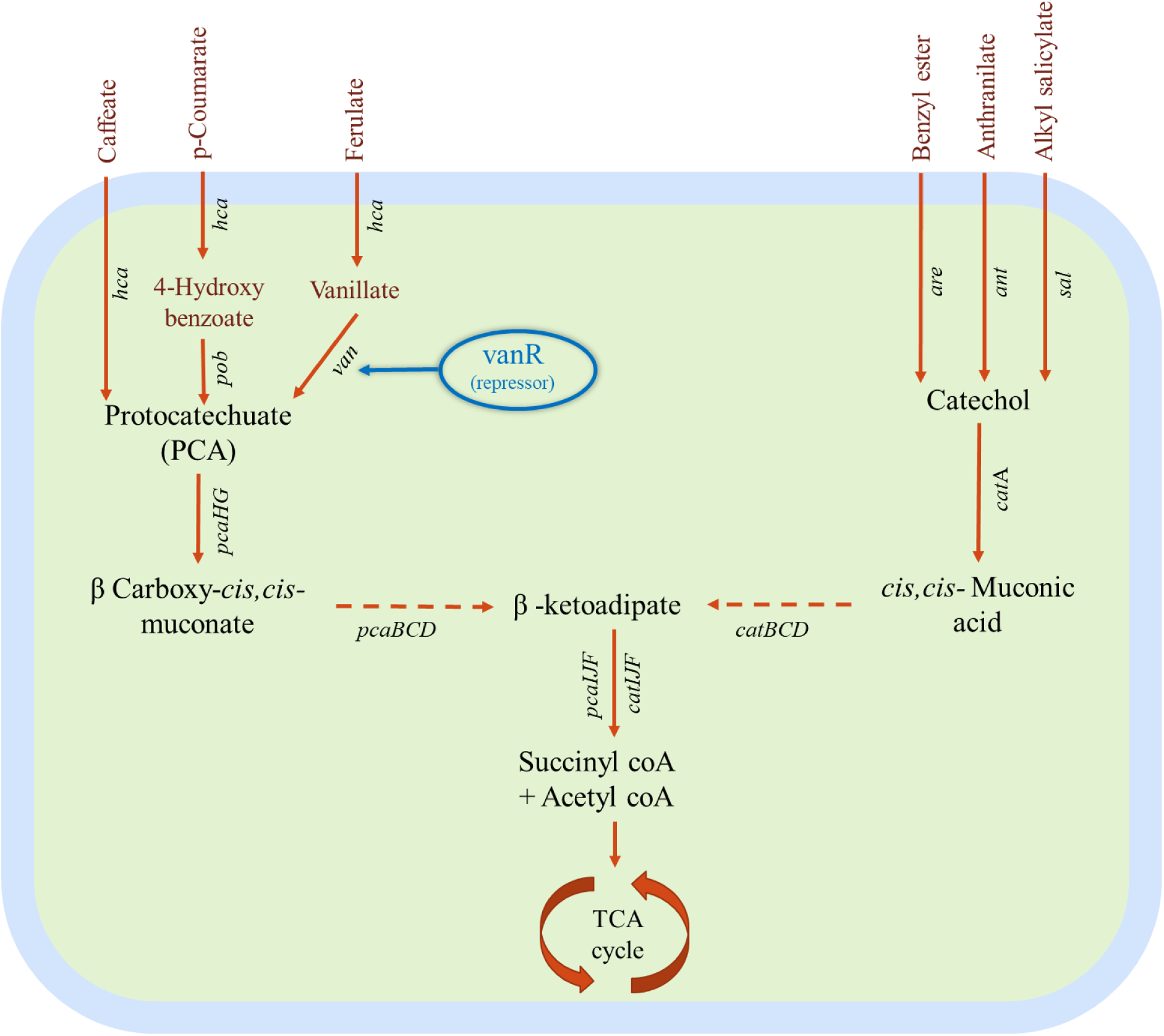
β-ketoadipate pathway of *Acinetobacter baylyi* ADP1 highlighting *vanR* gene., which acts as a repressor for *vanA*B genes.

### ALE of ADP1 strains on mixture of multiple LRAs

The strains evolved on a single LRA preferentially utilized only that LRA at a high concentration and not the other LRAs (Supplemental Material 1 Fig. S2). Therefore, we attempted to further evolve each of these strains to grow on a mixture of LRAs by sequentially adapting them to an additional LRA (Supplemental Material 1 Table. S3). Out of 20 strains that we tested, 3 showed stable growth and could simultaneously utilize 2 LRAs at higher concentrations, but not more (Supplemental Material 1 Fig. S3). Therefore, we attempted to grow all the evolved strains on MM-I, which contained a mixture of all five LRAs with a cumulative concentration of 60 mM (Table 2). We observed that none of the evolved strains could grow on MM-I (Supplemental Material 1 Fig. S4). We had observed earlier that wild-type ADP1 could grow on a medium containing minimal concentration of all the 5 monomers, with a cumulative concentration of 16 mM (Supplemental Material 1 Fig. S5a), but it also could not grow on MM-I (Supplemental Material 1 Fig. S5e).

Subsequently, a consortium of all the five single-LRA-evolved ADP1 strains, i.e., SAG_Cm, SAG_Hb, SAG_Fe, SAG_Vn and SAG_Cf, was grown (in quintuplicates) at different concentrations of the LRAs (Supplemental Material 1 Fig. S5b, d, f). After observing that three of these consortium cultures effectively consumed all the LRAs at low concentrations as well as in MM-I, we increased the concentrations of all the LRAs to that in the MM-II medium (Table 1). Through this process, we adapted the three consortia to the LRA- mixture in MM-II (Fig. 2). Subsequently, the concentration of some LRAs was further increased to that in the MM-III medium, while capping the total concentration to 100 mM (Table 1). However, we found that the consortia adapted to MM-II could not take up the LRAs in the MM-III medium. We plated the MM-II adapted consortium-culture on MM-II agar plates and cultured a few of the isolated colonies on MM-II liquid media. Through this process, we obtained a unique strain (SAG_185) which could grow on MM-II media and consume all the LRAs in this mixture (Fig. 3). However, SAG_185 also did not grow on MM-III.

**Fig. 2.**
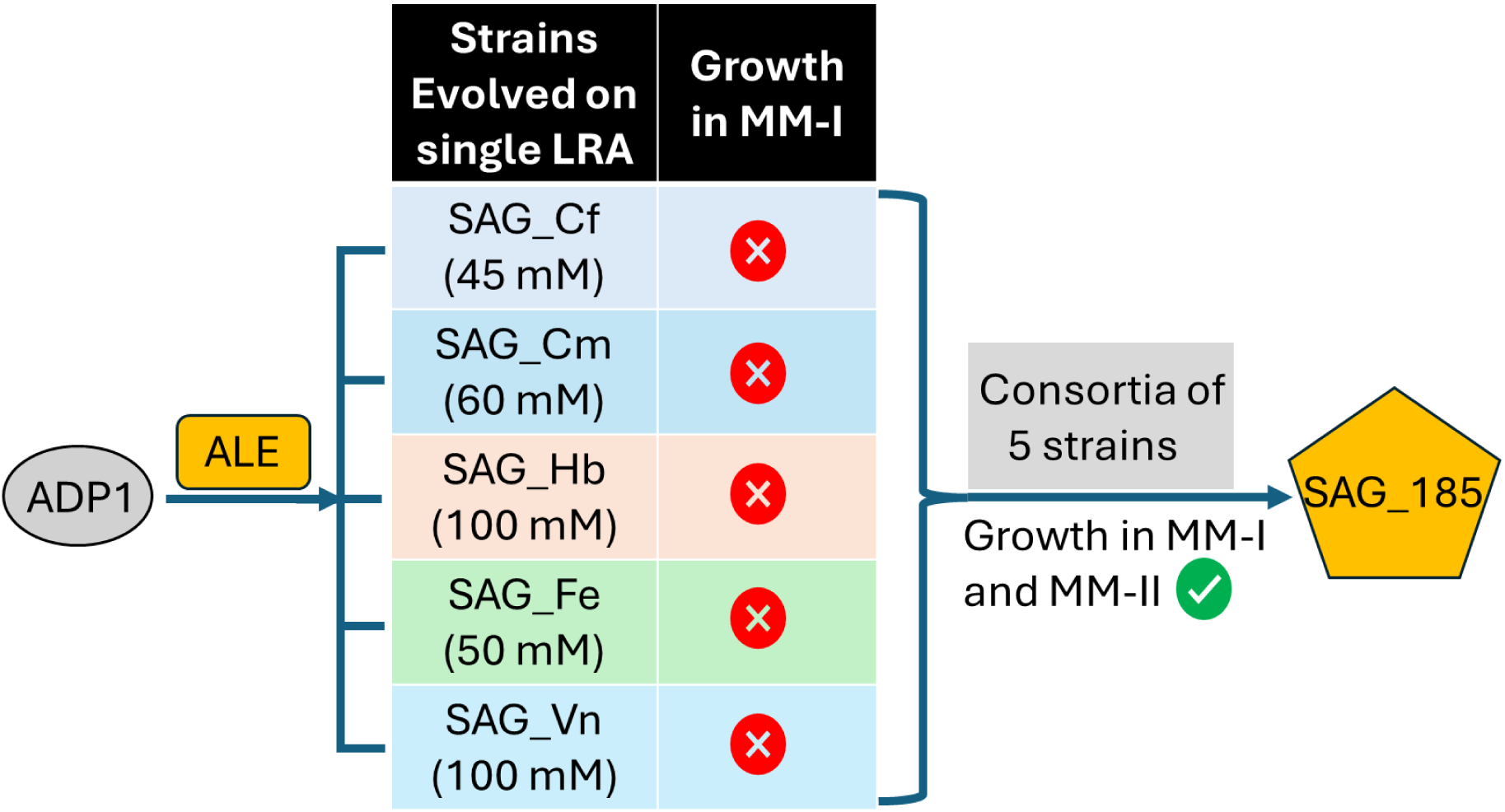
Overview of the ALE process of evolved-ADP1 consortia in MM-I and MM-II to obtain SAG_185.

**Fig. 3.**
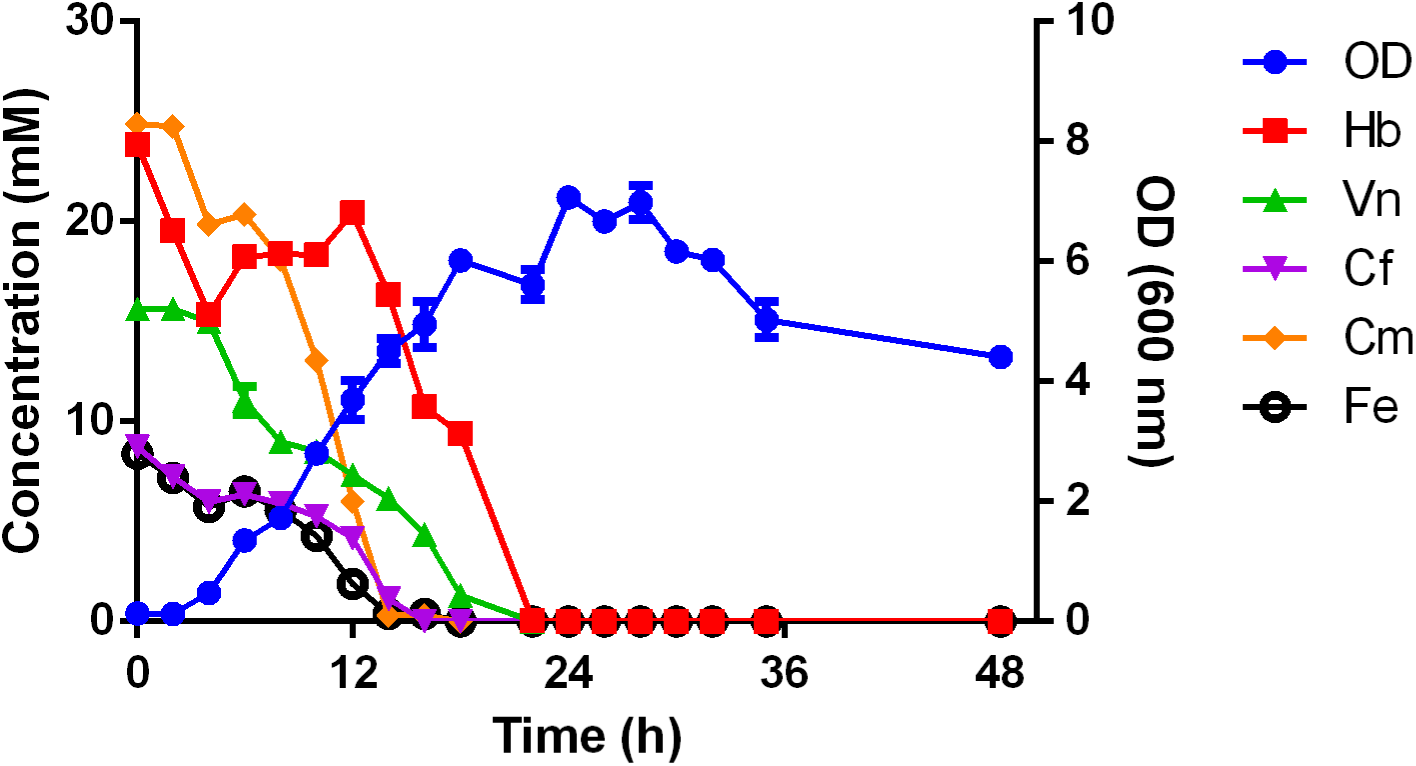
Growth and substrate utilization kinetics of SAG_185 in synthetic LRA mixture (MM-II). Cm: p- Coumaric acid, Hb: 4-hydroxybenzoic acid, Cf: Caffeic acid, Fe: Ferulic acid, Vn: Vanillic acid. OD denotes optical density at 600 nm.

The reproducibility of evolving a unique strain through the above consortium-evolution approach was validated by multiple experiments involving a consortium of all the five strains grown in MM-II medium. Single colonies isolated from these cultures exhibited similar growth and substrate utilization pattern. Additional data for one such strain, comparable to the profile for SAG_185 (Fig. 3), is shown in the Supplemental Material 1 Fig. S6.

We monitored the uptake profiles of all the 5 LRAs in the SAG_185 culture grown on MM-II media and observed that the culture consumed all the substrates simultaneously (Fig. 3). However, while the strain spontaneously consumed 4-hydroxybenzoic acid as the most preferred substrate, other substrates were consumed more slowly. We observed an accumulation of 4-hydroxybenzoic acid after 4th hour in the SAG_185 culture, which is due to the bioconversion of p-coumaric acid to 4-hydroxybenzoic acid (Fig. 1). 4- hydroxybenzoic acid was again consumed after the depletion of p-coumaric acid (Fig. 3). Since all the 5 LRAs from MM-II media were utilized by the SAG_185 culture, we also measured the levels of a key intermediate in the β-ketoadipate pathway, viz. PCA during the mid-log phase, in SAG_185 and a few other evolved strains. We found substantial accumulation of PCA in SAG_185 (Fig. 4), which could be useful for metabolic engineering of this strain.

**Fig. 4.**
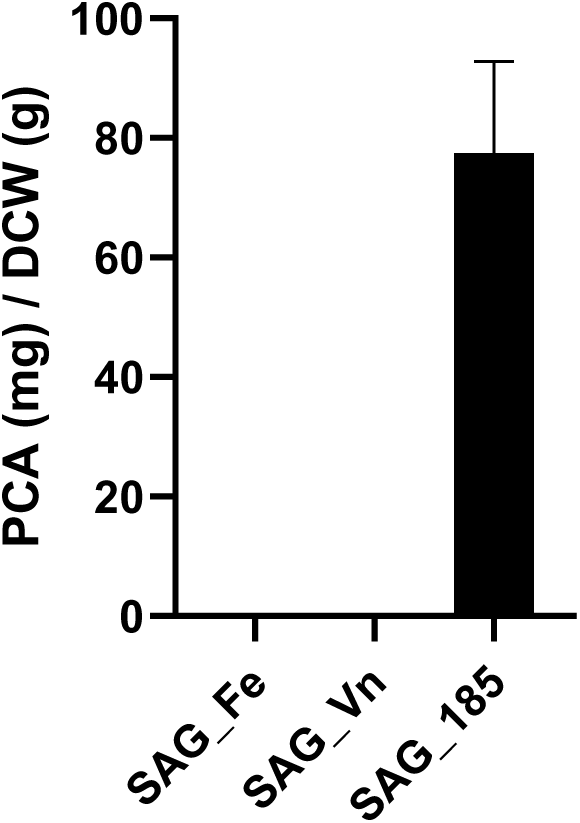
The levels of protocatechuate (PCA) during mid-log phase in evolved strains. No detectable levels of PCA were present in SAG_Fe and SAG_Vn. Error bars represent SD (n=2)

### Growth studies on depolymerized lignin

A lignin-rich biomass, obtained after cellulosic hydrolysis of pre-treated corncob, was depolymerized by base-catalysed depolymerization (BCD). The depolymerized lignin was found to predominantly contain p-coumaric acid (36.66 ± 4.09 mM) and minimal concentrations of ferulic acid (4.82 ± 0.59 mM). Other aromatic monomers and oligomers were not present in detectable levels in the mixture. The wild-type ADP1 did not grow properly on minimal media supplemented with depolymerized lignin (BCD_Cm20) or utilize the p-coumaric acid. Since SAG_Cm was adapted only in p-coumaric acid, the growth of SAG_Cm on depolymerized lignin was compared with that of SAG_185. Although the adaptive evolution of SAG_Cm allowed for growth in 60 mM synthetic p-coumaric acid, this strain was still unable to utilize or grow on BCD_Cm20 (Fig. 5). This may be due to inhibition by other LRAs or inhibitory compounds in the depolymerized lignin. However, SAG_185 was able to grow and utilize p-coumaric acid from the BCD_Cm20 media, with intermediate production and consumption of 4-hydroxybenzoic acid (Fig. 6).

**Fig. 5.**
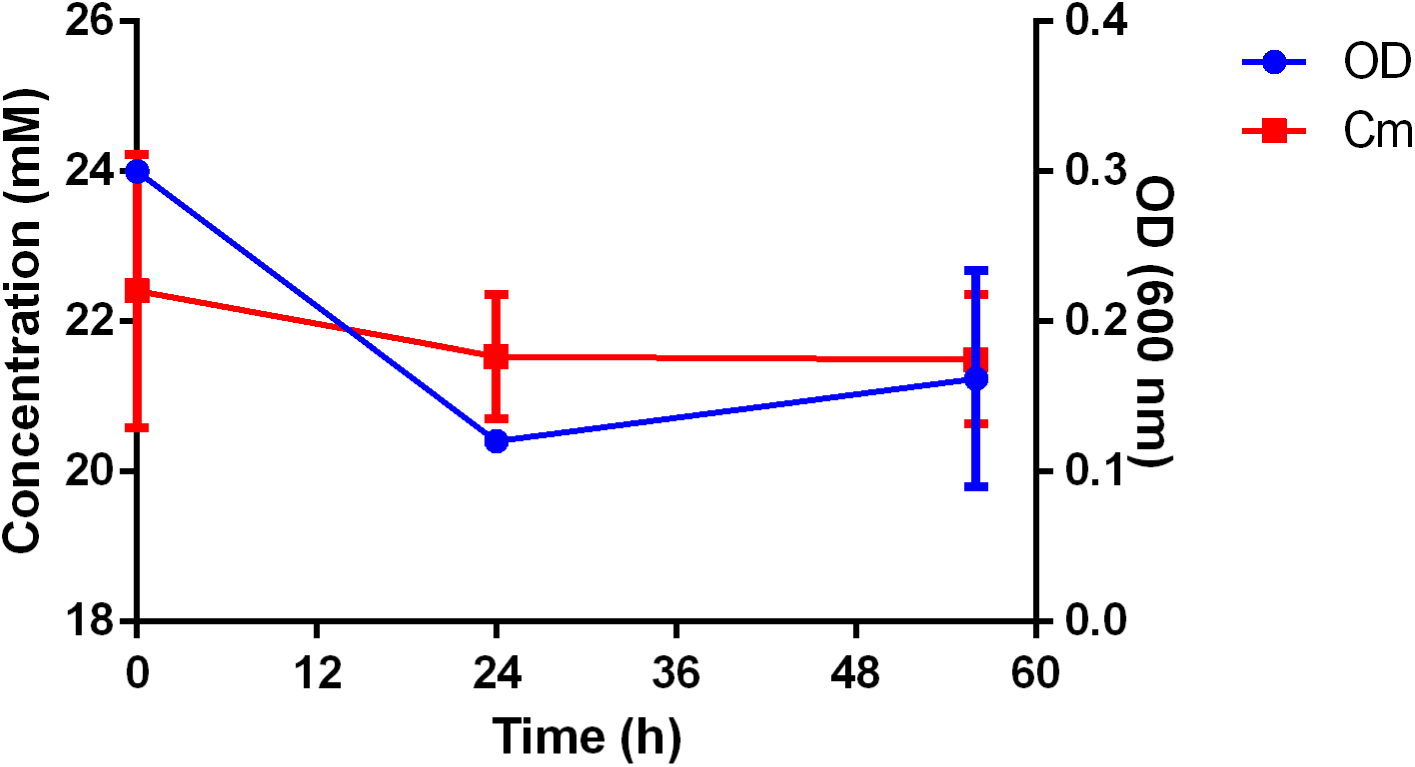
Growth and substrate utilization of SAG_Cm on depolymerized lignin.

**Fig. 6.**
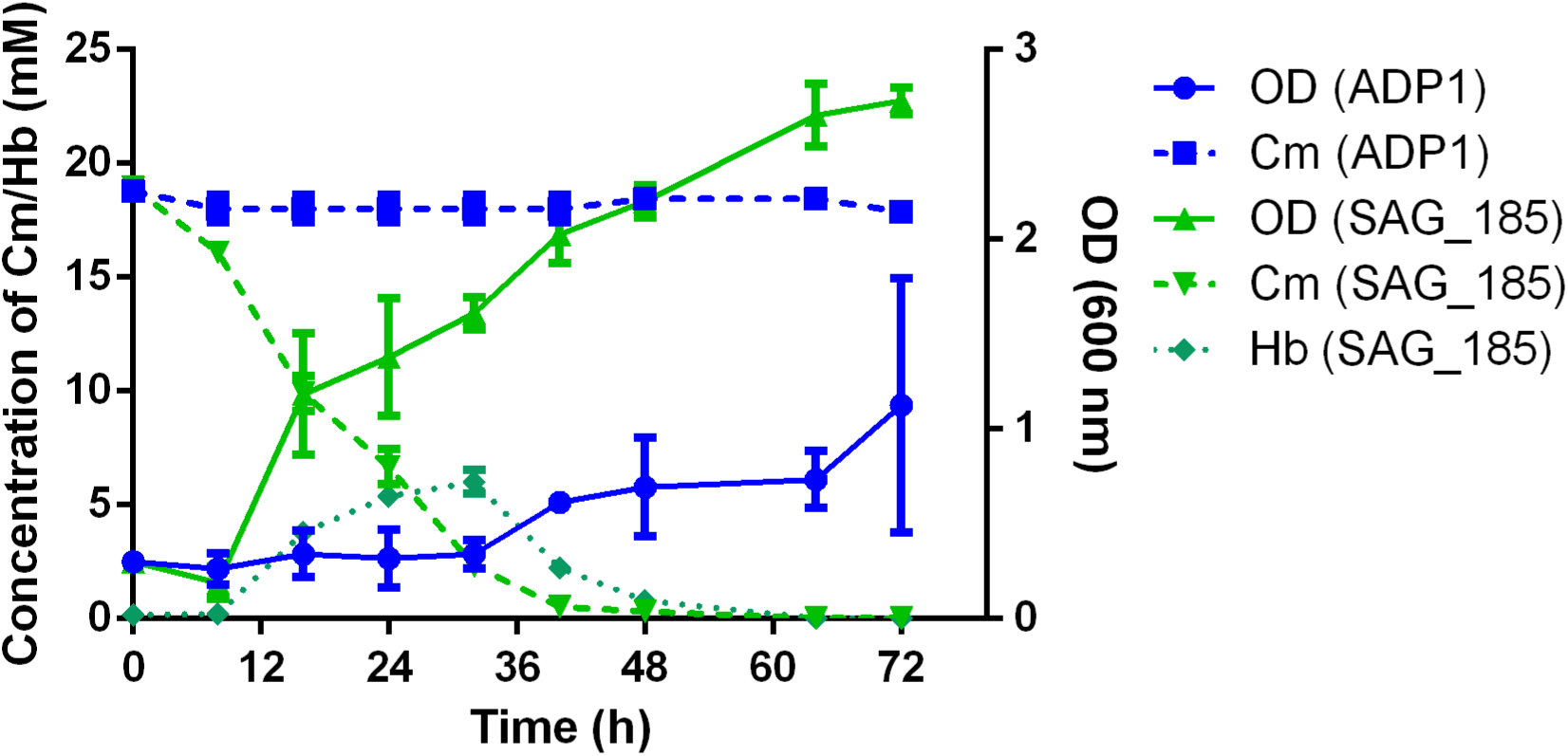
Growth comparison of ADP1 and SAG_185 on BCD_Cm20 media and comparison of p-coumaric acid (Cm) utilization by ADP1 and SAG_185. Increase in 4-hydrozybenzoic acid (Hb) concentration is due to bioconversion of Cm to its intermediate, Hb. Hb is consumed after the exhaustion of Cm.

A previous study has shown that the growth of *Pseudomonas putida* KT2440 was affected when grown on depolymerized lignin (38). In this study, we observed that evolving the strain simultaneously in multiple LRAs results in the overcoming of this inhibition. To investigate the genetic modifications leading to these phenotypic changes, we carried out WGS and comparative analyses of all the evolved ADP1 strains.

### Identification of potential targets by genome sequencing of evolved strains

All the independently evolved strains were genome-sequenced to identify the genetic changes which could explain the adaptation to higher concentration of the LRAs and growth on depolymerized lignin. In this study, the wild-type ADP1 served as the reference genome for identifying all the mutations. The comparison between wild-type ADP1 and the reference ADP1 genome (NCBI GenBank: CR543861.1) revealed a total of 14 mutational differences (Supplemental Material 2 Table S1), which were consistent with a previous study (39).

The sequence data revealed that all the evolved strains had very strong sequence similarity with ADP1 (Supplemental Material 1 Table. S4). All the evolved strains, except SAG_Hb, exhibited several mutations (Fig. 7, Supplemental Material 1 Fig. S7 and, Supplemental Material 2 Table S1 and S2). The genetic changes in the evolved strains involved various types of mutations, including IS-mediated insertions, deletions, and base substitutions (Supplemental Material 1 Fig. S7). Most of the genetic changes (∼ 57% of the mutations) observed in all the evolved strains were induced due to IS1236 insertion sequences (Supplemental Material 2 Table S1 and S2), with similar observations in earlier literature reports (27). The IS1236 insertion sequences are involved in the duplication of a 3-bp DNA segment flanking the IS element insertion site (40).

**Fig. 7.**
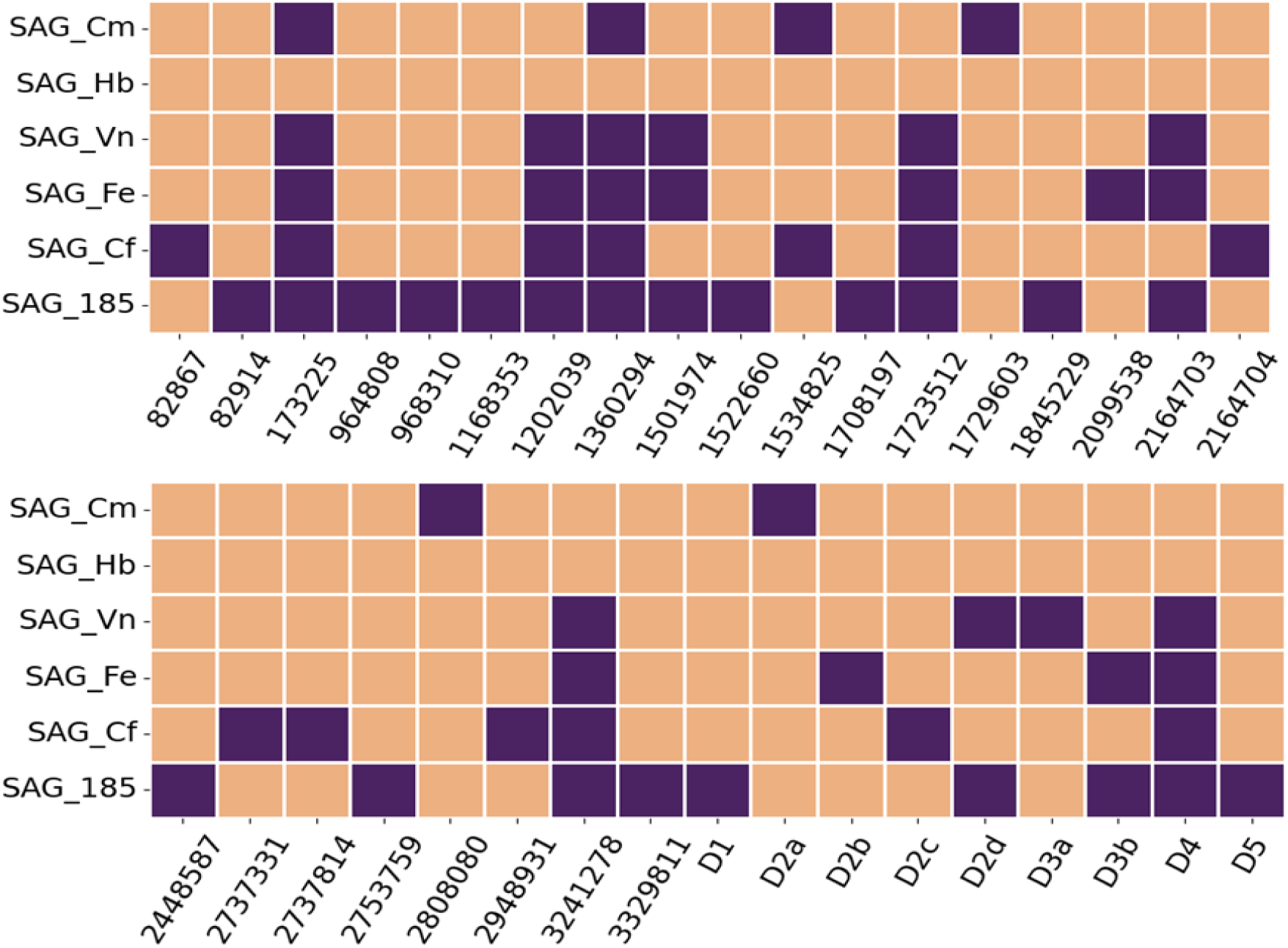
Comparative analysis of the genetic modifications in the individual ADP1 evolved strains and SAG_185. The x-axis denotes the base position number corresponding to the reference genome (GenBank CR543861.1). D1-D5 denotes large deletion regions (Supplemental Material 2 Table S2). The violet color indicates mutation at that base position.

We observed 22 mutations in SAG_185, of which 12 mutations were unique, i.e. not present in any of the other strains evolved on single LRAs. These mutations probably occurred when the consortium of strains was grown and evolved on the mixture of LRAs. The other 10 mutations were shared with the different strains evolved on single LRAs. Specifically, SAG_Cm had two common mutations, SAG_Cf had six common mutations, and SAG_Vn and SAG_Fe had nine common mutations each with SAG_185. This indicated that SAG_185 probably evolved from either the SAG_Vn or SAG_Fe strain (Fig. 7).

Two of the unique mutations in SAG_185, *vanR* and *pcaU*, were in important regulatory genes. These genes directly influence the uptake of lignin monomers through the β-ketoadipate pathway (Fig. 1 and Supplemental Material 1 Table S5) (1). VanR acts as a repressor controlling the expression of *vanAB* genes (41), whereas PcaU acts as a bifunctional transcriptional regulator of the structural genes present in the *pca* operon, with the functionality depending on the level of PCA (42) PcaU acts as an activator for the *pca* operon in the presence of elevated PCA levels and as a repressor when PCA concentration is low, for its own expression as well as the structural genes in the *pca* operon. The *pca* operon consists of nine genes (*pcaIJFBDKCHG*) that play a crucial role in the catabolism of PCA, converging into the β-ketoadipate pathway at β-ketoadipate-CoA and ultimately entering the TCA cycle as acetyl-CoA (1). Luo et al. (2022) (27) have reported an SNP in *pcaU* due to ALE of ADP1 on ferulate, which was identical to the one found in our whole-genome sequence. However, Sanger-sequencing of the *pcaU* gene in SAG_185 showed no SNP present in the gene.

### Gene expression studies in β-ketoadipate pathway using RT-qPCR

Based on the WGS analysis, we conducted RT-qPCR studies on three specific genes (*vanA*, *vanB* and *vanR*) that are involved in the β-ketoadipate pathway. VanR, belonging to the GntR repressor family of proteins (41), represses *vanA* and *vanB* genes coding for vanillate demethylase, which converts vanillate to PCA (43). We observed increased expression of *vanA* and *vanB* genes in the SAG_Fe, SAG_Vn, and SAG_185 strains (P < 0.001), compared to ADP1 (Fig. 8). A lower expression of *vanR* in SAG_Fe resulted in higher levels of *vanAB*.

**Fig. 8.**
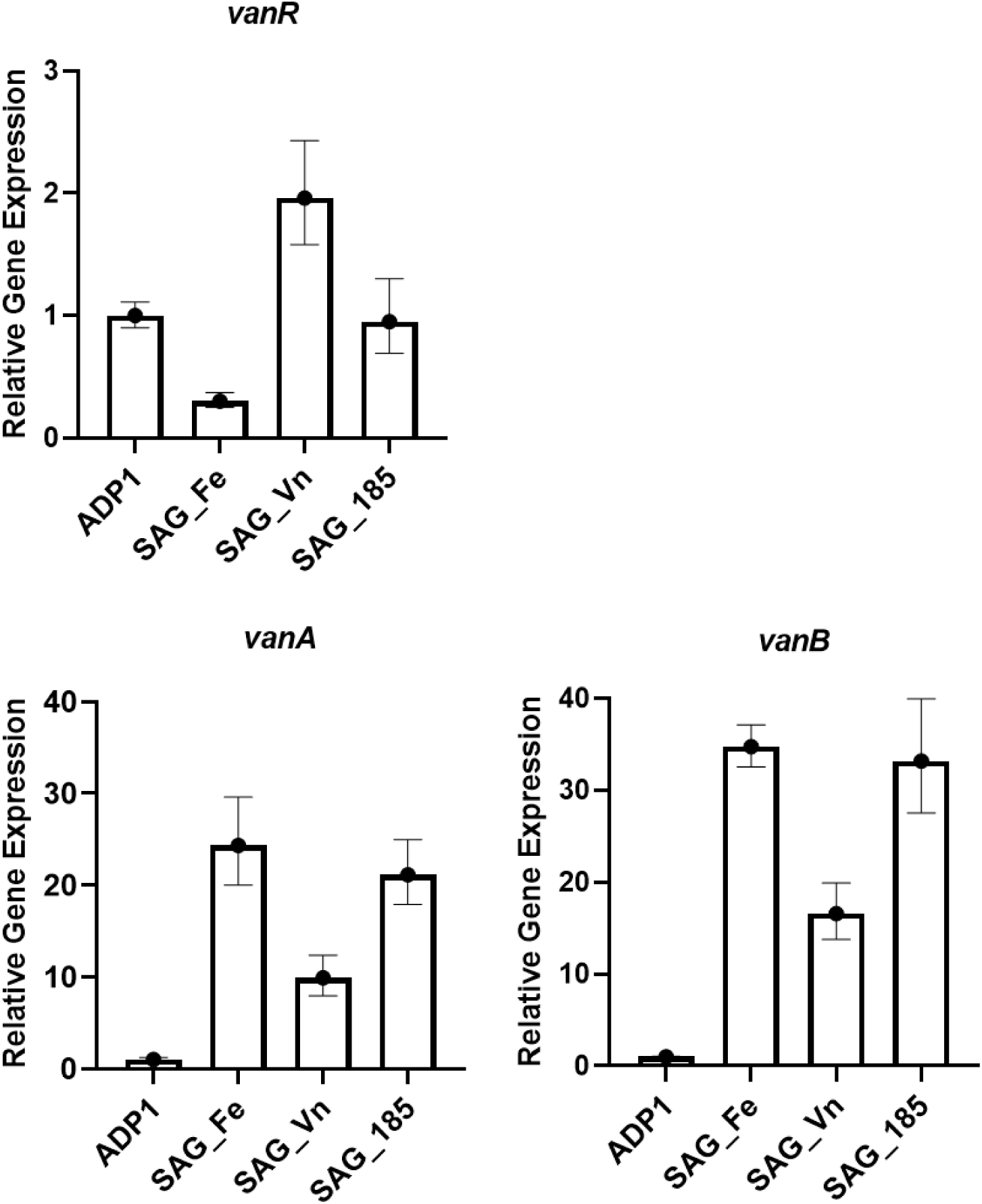
Relative Gene Expression (RGE) of *vanA*, *vanB*, and *vanR* in ADP1, SAG_Fe, SAG_Vn, and SAG_185. The data is normalized with respect to wild-type ADP1 (RGE = 1). The RT-qPCR experiment was done with three biological replicates each having two technical replicates. The error bar represents the standard error.

We observed a higher level of *vanR* expression in SAG_185 relative to SAG_Fe. However, the *vanR* mutation in SAG_185 probably resulted in an increased expression of *vanAB* genes in this strain, comparable to SAG_Fe (P > 0.05). In SAG_Vn, despite the high expression of *vanR*, we observed a reasonably enhanced expression of the *vanAB* genes in this strain, possibly due to the high levels of vanillate in the culture medium. A previous study reported a basal level of *vanAB* expression in the absence of vanillate. However, both vanillate-induced ADP1 and Δ*vanR* ADP1 mutant strains show increased expression of *vanAB* genes (41). This indicates the role of vanillate as an inducer and *vanR* in transcriptional repression of *vanAB* genes.

### Other genetic changes in the evolved strains based on WGS

Overall, 35 genetic changes were present in all the 6 evolved strains and 16 of these were present in the genes coding for either a putative/hypothetical protein or found in the intergenic region (Supplemental Material 2 Table S1 and S2). There were 9 large deletion regions (>172 nt) (Supplemental Material 2 Table S2) out of which 1 region lies in the intergenic region and 6 regions mostly encode for the hypothetical/putative proteins. These include IS-element affected coding regions such as *ACIAD1523* (ATP-binding cassette transporter), which is known to be involved in the uptake of carbohydrates (44). In addition, the IS-element insertion in the gene for the competence factor, *comP* (*ACIAD3338*), involved in the uptake of DNA (45), led to the loss of natural transformation in the evolved strains.

Additionally, there were two large deletions comprising of 9kb and 38kb regions. In the 9 kb deletion region, most genes were hypothetical/putative proteins, and the rest belonged to the IS family. In the 38kb deletion region, the sequences coding for putative/hypothetical proteins included transcriptional regulators of LysR family (*ACIAD1543*), MarR family (*ACIAD1559*) and AraC family (*ACIAD1576*). These family of regulators play a role in the degradation of p-coumarate, catechol and benzoate in *Acinetobacter*; p-hydroxybenzoate and benzoate in *Pseudomonas* (46, 47). However, there is no direct evidence of the role of these three genes in the regulation of the β-ketoadipate pathway, even though the family of transcriptional regulators is the same. Additionally, other genes in this 38kb deletion region code for the utilization of tricaballylate (*tcuC*, *tcuA*, *tcuR*) (48) and fatty acid metabolism (*ACIAD1554*, *ACIAD1565*, *ACIAD1566*, *ACIAD1568*, *ACIAD1569*, *ACIAD1570*, *ACIAD1572*), as well as the degradation of coniferyl alcohol (*ACIAD1578 xylB*). A previous study shows that the degradation of coniferyl alcohol reduced significantly in the ADP1Δ*xylB* strain but did not stop completely (49).

In SAG_185, apart from mutation in *vanR*, there were mutations in *quiA* and *hcaE* genes which are also part of the β-ketoadipate pathway (Supplemental Material 2 Table S1 and S2). The gene *quiA* encodes quinate/shikimate dehydrogenase, an enzyme that converts quinate and shikimate into dehydroquinate and dehydroshikimate, respectively. The β-ketoadipate pathway further processes these compounds into PCA (50). Previous ALE studies on ADP1 using ferulate as a substrate (27) revealed mutations in the aromatic transporter genes, such as *hcaE*, *hcaK*, and *vanK*, and the gene associated with lipopolysaccharide synthesis (ACIAD0482). The *hcaE* gene encodes a porin that belongs to the OprD family and shares significant similarity with the porin-encoding gene *vanP*, which is involved in vanillate transport (51). The *hcaE* gene belongs to the *hca* operon consisting of five genes: *hcaABCDE* which helps in the catabolism of LRAs such as caffeate, coumarate and ferulate (52). In our study, all the evolved strains, except SAG_Hb, had mutations in the *hcaE* gene. WGS analysis of SAG_Fe, SAG_Vn, SAG_Cf and SAG_185 revealed a deletion of 207 nucleotides within the *hca* operon, specifically covering 184 nucleotides from the downstream end of the *hcaE* gene (Supplemental Material 2 Table S2). A previous study by Luo et al. (2022) (27) revealed that mutations in the *hcaE* was beneficial for the growth of *A. baylyi* ADP1 evolved on ferulate, and this beneficial effect was confirmed by reverse engineering. They also reported mutations in *ACIAD3465* (putative two-component sensor) and *ACIAD2867* (putative H^+^/Na^+^ antiporter). We found similar mutations in our study but at different positions within these genes.

## DISCUSSION

Previous literature reports on ALE of strains involved in utilization of LRA have focused only on improving the strain tolerance to a single monomer. However, depolymerization of lignin results in multiple monomers which need to be utilized. For example, the depolymerized lignin fraction obtained in this study predominantly contains p-coumaric acid, small amounts of ferulic acid and minimal amounts of other monomers/oligomers. Hence, we proceeded to evolve a strain tolerant to presence of multiple monomers in the medium. Development of an efficient fermentation process also requires substrates at high concentrations. Although wild-type ADP1 can withstand multiple monomers at minimal concentrations, it was difficult to adapt it and make it grow at higher concentration of LRAs (Supplemental Material 1 Fig S5 a,c,e).

In this work, we have demonstrated a two-stage approach to ALE for achieving the above objectives. Initially, ADP1 was evolved on single synthetic LRAs and five evolved strains were obtained which were respectively adapted to a high concentration of 5 different LRAs (Table 2). It was attempted to evolve each of these strains to utilize multiple LRAs, which was only partially successful since only one additional LRA was taken up (Supplemental Material 1 Fig S3, Table S3). In a subsequent approach, a consortium of the five evolved strains were grown on a mixture of the 5 LRAs (MM-I). Since the consortium successfully consumed all the LRAs, it was subjected to ALE by increasing the concentration of all the LRAs (MM-II). The adaptive evolution of the consortium resulted in a unique strain, SAG_185, with the ability to simultaneously utilize all the LRAs in MM-II. There are no literature reports available on the consortium-evolution approach that we have described in this study. This novel ALE process could therefore be adapted for evolution of other strains which are required to grow on a mixture of toxic substrates.

In addition to simultaneous uptake of LRAs in the synthetic mixture, SAG_185 was also able to grow on depolymerized lignin medium (BCD_Cm20), which contained p-coumarate (20mM) as the predominant lignin-derived monomer (LDM) and low concentrations of other LDMs. In contrast, none of the strains which were evolved on individual synthetic LRAs were able to grow on depolymerized lignin. To understand the differences in the physiology between SAG_185, the other evolved strains and the wild-type ADP1, we subjected them to WGS and performed a comparative analysis.

Most of the genetic changes were observed in genes which code for hypothetical proteins, transporters and putative transcriptional regulators. Majority of them were changes due to IS1236 insertion sequences, where IS1236 is involved in the duplication of a 3-bp DNA segment flanking the IS element insertion site (40). The role of these hypothetical protein regions and regulators in the uptake of LRAs is still unknown, indicating that there are far more genetic regions affecting the uptake of these aromatics.

WGS analysis revealed 10 mutations in SAG_185 identical to the ones in SAG_Vn and SAG_Fe (Fig. 7). ADP1 is known for its natural competence and high frequency of homologous recombination (53–55). Previous studies have reported intra and inter-species horizontal gene transfers in *Acinetobacter* sps. (56, 57). Therefore, it is possible that one of the evolved strains underwent further genetic changes when the consortium was grown on a mixture of LRAs and acquired beneficial traits to grow in the LRA mixture.

Although there were 12 unique mutations present in SAG_185, the most interesting finding in our study is the mutation in *vanR* gene, which is directly involved in the catabolism of aromatic compounds through the β-ketoadipate pathway (1). Deletion of the VanR binding site in the promoter region of *vanABK* operon of *Corneybacterium glutamicum* resulted in constitutive expression of *vanABK* operon (58). The IS-mediated mutation of *vanR* in SAG_185 may have resulted in a non-functional protein which was unable to repress *vanAB* expression in SAG_185.

PCA is a key common intermediate of the β-ketoadipate pathway in ADP1, arising due to the consumption of each of the LRAs taken up in this study (Fig. 1). We found substantial accumulation of PCA in the SAG_185 culture, which was not detectable in SAG_Vn and SAG_Fe cultures. Further bioconversion of PCA to other downstream metabolites, including β-ketoadipate, is tightly regulated through the *pca* operon and therefore this could be a potential target for metabolic engineering. PCA has important applications in biomedicine (59), food preservatives (60) and as a building block of biodegradable polyesters (61). The insertion elements play a role in gene duplication and amplification events (62) and their removal enhances genome stability in ADP1 (39). Deletion mutants (Δ*IS1236*) of these insertion sequences in ADP1 have demonstrated reduced mutation rates and improved transformation efficiency (63). Thus, utilising the ADP1Δ*IS1236* for strain engineering would be advantageous for achieving long-term stable expression of heterologous genes. The insights obtained from this work will be used for developing a platform *A. baylyi* strain by rational metabolic engineering.

## Data availability

All the sequencing data were deposited in the National Center for Biotechnology Information (NCBI) database under BioProject accession number PRJNA1027358 (https://www.ncbi.nlm.nih.gov/sra/PRJNA1027358).

## Contributions

ABM and GJ conceptualized the project; SM, PS, ABM contributed to the experimental design; SM performed the ALE experiments, analytics and growth studies on depolymerized lignin; PS performed the WGS analysis, RT-qPCR experiments and statistical analysis; VPJ contributed to lignin de-polymerization by BCD; SM, PS and GJ wrote the manuscript; GJ was the Principal Investigator for this project and recipient of the project funding; ABM and LMB were the co-Principal Investigators for this project. All authors read and approved the manuscript.

## Ethics declarations

### Human and animal rights

This article does not contain any studies with human participants or animals performed by any of the authors.

## References

1. Harwood CS, Parales RE. 1996. The beta-ketoadipate pathway and the biology of self-identity. Annu Rev Microbiol 50:553–590.

2. Young DM, Parke D, Ornston LN. 2005. Opportunities for genetic investigation afforded by *Acinetobacter baylyi*, a nutritionally versatile bacterial species that is highly competent for natural transformation. Annu Rev Microbiol 59:519–551.

3. Biggs BW, Bedore SR, Arvay E, Huang S, Subramanian H, McIntyre EA, Duscent-Maitland C V., Neidle EL, Tyo KEJ. 2020. Development of a genetic toolset for the highly engineerable and metabolically versatile *Acinetobacter baylyi* ADP1. Nucleic Acids Res 48:5169–5182.

4. Suárez GA, Dugan KR, Renda BA, Leonard SP, Gangavarapu LS, Barrick JE. 2021. Rapid and assured genetic engineering methods applied to *Acinetobacter baylyi* ADP1 genome streamlining. Nucleic Acids Res 48:4585–4600.

5. Elbahloul Y, Steinbüchel A. 2006. Engineering the genotype of *Acinetobacter* sp. strain ADP1 to enhance biosynthesis of cyanophycin. Appl Environ Microbiol 72:1410–1419.

6. Salmela M, Lehtinen T, Efimova E, Santala S, Santala V. 2019. Alkane and wax ester production from lignin-related aromatic compounds. Biotechnol Bioeng 116:1934–1945.

7. Luo J, Efimova E, Losoi P, Santala V, Santala S. 2020. Wax ester production in nitrogen-rich conditions by metabolically engineered *Acinetobacter baylyi* ADP1. Metab Eng Commun 10:e00128.

8. Santala S, Efimova E, Kivinen V, Larjo A, Aho T, Karp M, Santala V. 2011. Improved Triacylglycerol Production in *Acinetobacter baylyi* ADP1 by Metabolic Engineering. Microbial Cell Factories 2011 10:1 10:1–10.

9. Luo J, Lehtinen T, Efimova E, Santala V, Santala S. 2019. Synthetic metabolic pathway for the production of 1-alkenes from lignin-derived molecules. Microbial Cell Factories 2019 18:1 18:1–13.

10. Salmela M, Lehtinen T, Efimova E, Santala S, Santala V. 2020. Towards bioproduction of poly-α-olefins from lignocellulose. Green Chemistry 22:5067–5076.

11. Arvay E, Biggs BW, Guerrero L, Jiang V, Tyo K. 2021. Engineering *Acinetobacter baylyi* ADP1 for mevalonate production from lignin-derived aromatic compounds. Metab Eng Commun 13:e00173.

12. Kurnia K, Efimova E, Santala V, Santala S. 2024. Metabolic engineering of *Acinetobacter baylyi* ADP1 for naringenin production. bioRxiv 2024.06.06.597799.

13. Adriaens P, Focht DD. 1991. Cometabolism of 3,4-dichlorobenzoate by *Acinetobacter* sp. strain 4-CB1. Appl Environ Microbiol 57:173–179.

14. Abdel-El-Haleem D. 2004. *Acinetobacter*: environmental and biotechnological applications. Afr J Biotechnol 2:71–74.

15. Delneri D, Degrassi G, Rizzo R, Bruschi C V. 1995. Degradation of trans-ferulic and p-coumaric acid by *Acinetobacter calcoaceticus* DSM 586. Biochimica et Biophysica Acta (BBA) -General Subjects 1244:363–367.

16. Johnson CW, Salvachúa D, Khanna P, Smith H, Peterson DJ, Beckham GT. 2016. Enhancing muconic acid production from glucose and lignin-derived aromatic compounds via increased protocatechuate decarboxylase activity. Metab Eng Commun 3:111–119.

17. Upadhyay P, Lali A. 2021. Protocatechuic acid production from lignin-associated phenolics. Prep Biochem Biotechnol 51:979–984.

18. Salvachúa D, Karp EM, Nimlos CT, Vardon DR, Beckham GT. 2015. Towards lignin consolidated bioprocessing: simultaneous lignin depolymerization and product generation by bacteria. Green Chemistry 17:4951–4967.

19. Min D, Xiang Z, Liu J, Jameel H, Chiang V, Jin Y, Chang H. 2015. Improved Protocol for Alkaline Nitrobenzene Oxidation of Woody and Non-Woody Biomass. Journal of Wood Chemistry and Technology 35:52–61.

20. Zhang X, Zhu J, Sun L, Yuan Q, Cheng G, Argyropoulos DS. 2019. Extraction and characterization of lignin from corncob residue after acid-catalyzed steam explosion pretreatment. Ind Crops Prod 133:241– 249.

21. Timokhin VI, Regner M, Motagamwala AH, Sener C, Karlen SD, Dumesic JA, Ralph J. 2020. Production of p-Coumaric Acid from Corn GVL-Lignin. ACS Sustain Chem Eng 8:17427–17438.

22. Kurosawa K, Laser J, Sinskey AJ. 2015. Tolerance and adaptive evolution of triacylglycerol-producing *Rhodococcus opacus* to lignocellulose-derived inhibitors. Biotechnology for Biofuels 2015 8:1 8:1–14.

23. Cerisy T, Souterre T, Torres-Romero I, Boutard M, Dubois I, Patrouix J, Labadie K, Berrabah W, Salanoubat M, Doring V, Tolonen AC. 2017. Evolution of a biomass-fermenting bacterium to resist lignin phenolics. Appl Environ Microbiol 83.

24. Wang X, Khushk I, Xiao Y, Gao Q, Bao J. 2018. Tolerance improvement of *Corynebacterium glutamicum* on lignocellulose derived inhibitors by adaptive evolution. Appl Microbiol Biotechnol 102:377–388.

25. Wang Q, Qi W, Wang W, Zhang Y, Leksawasdi N, Zhuang X, Yu Q, Yuan Z. 2019. Production of furfural with high yields from corncob under extremely low water/solid ratios. Renew Energy 144:139– 146.

26. Pardo I, Jha RK, Bermel RE, Bratti F, Gaddis M, McIntyre E, Michener W, Neidle EL, Dale T, Beckham GT, Johnson CW. 2020. Gene amplification, laboratory evolution, and biosensor screening reveal MucK as a terephthalic acid transporter in *Acinetobacter baylyi* ADP1. Metab Eng 62:260–274.

27. Luo J, McIntyre EA, Bedore SR, Santala V, Neidle EL, Santala S. 2022. Characterization of Highly Ferulate-Tolerant *Acinetobacter baylyi* ADP1 Isolates by a Rapid Reverse Engineering Method. Appl Environ Microbiol 88.

28. Mohamed ET, Werner AZ, Salvachúa D, Singer CA, Szostkiewicz K, Rafael Jiménez-Díaz M, Eng T, Radi MS, Simmons BA, Mukhopadhyay A, Herrgård MJ, Singer SW, Beckham GT, Feist AM. 2020. Adaptive laboratory evolution of Pseudomonas putida KT2440 improves p-coumaric and ferulic acid catabolism and tolerance. Metab Eng Commun 11:e00143.

29. Kannisto M, Efimova E, Karp M, Santala V. 2017. Growth and wax ester production of an *Acinetobacter baylyi* ADP1 mutant deficient in exopolysaccharide capsule synthesis. J Ind Microbiol Biotechnol 44:99–105.

30. Kannisto MS, Mangayil RK, Shrivastava-Bhattacharya A, Pletschke BI, Karp MT, Santala VP. 2015. Metabolic engineering of *Acinetobacter baylyi* ADP1 for removal of *Clostridium butyricum* growth inhibitors produced from lignocellulosic hydrolysates. Biotechnology for Biofuels 2015 8:1 8:1–10.

31. Salmela M, Sanmark H, Efimova E, Efimov A, Hytönen VP, Lamminmäki U, Santala S, Santala V. 2018. Molecular tools for selective recovery and detection of lignin-derived molecules. Green Chemistry 20:2829–2839.

32. Schwarz M, Rodríguez MC, Guillén DA, Barroso CG. 2009. Development and validation of UPLC for the determination of phenolic compounds and furanic derivatives in Brandy de Jerez. J Sep Sci 32:1782–1790.

33. Deatherage DE, Barrick JE. 2014. Identification of mutations in laboratory-evolved microbes from next-generation sequencing data using breseq. Methods Mol Biol 1151:165–188.

34. Li H. 2013. Aligning sequence reads, clone sequences and assembly contigs with BWA-MEM. arXiv preprint arXiv:1303.3997

35. Li H, Handsaker B, Wysoker A, Fennell T, Ruan J, Homer N, Marth G, Abecasis G, Durbin R, Subgroup 1000 Genome Project Data Processing. 2009. The Sequence Alignment/Map format and SAMtools. Bioinformatics 25:2078–2079.

36. Svec D, Tichopad A, Novosadova V, Pfaffl MW, Kubista M. 2015. How good is a PCR efficiency estimate: Recommendations for precise and robust qPCR efficiency assessments. Biomol Detect Quantif 3:9–16.

37. Pfaffl MW. 2001. A new mathematical model for relative quantification in real-time RT–PCR. Nucleic Acids Res 29:e45–e45.

38. Almqvist H, Veras H, Li K, Garcia Hidalgo J, Hulteberg C, Gorwa-Grauslund M, Skorupa Parachin N, Carlquist M. 2021. Muconic Acid Production Using Engineered *Pseudomonas putida* KT2440 and a Guaiacol-Rich Fraction Derived from Kraft Lignin. ACS Sustain Chem Eng 9:8097–8106.

39. Renda BA, Dasgupta A, Leon D, Barrick JE. 2015. Genome instability mediates the loss of key traits by *Acinetobacter baylyi* ADP1 during laboratory evolution. J Bacteriol 197:872–881.

40. Gerischer U, D’Argenio DA, Ornston LN. 1996. IS 1236, a newly discovered member of the IS3 family, exhibits varied patterns of insertion into the *Acinetobacter calcoaceticus* chromosome. Microbiology (N Y) 142:1825–1831.

41. Morawski B, Segura A, Ornston LN. 2000. Repression of *Acinetobacter* vanillate demethylase synthesis by VanR, a member of the GntR family of transcriptional regulators. FEMS Microbiol Lett 187:65–68.

42. Trautwein G, Gerischer U. 2001. Effects exerted by transcriptional regulator PcaU from *Acinetobacter* sp. strain ADP1. J Bacteriol 183:873–881.

43. Segura A, Bünz P V., D’Argenio DA, Ornston LN. 1999. Genetic analysis of a chromosomal region containing *vanA* and *vanB*, genes required for conversion of either ferulate or vanillate to protocatechuate in *Acinetobacter*. J Bacteriol 181:3494–3504.

44. Saier MH. 2000. Families of transmembrane sugar transport proteins. Mol Microbiol 35:699–710.

45. Porstendörfer D, Gohl O, Mayer F, Averhoff B. 2000. ComP, a pilin-like protein essential for natural competence in *Acinetobacter* sp. strain BD413: Regulation, modification, and cellular localization. J Bacteriol 182:3673–3680.

46. Grove A. 2017. Regulation of Metabolic Pathways by MarR Family Transcription Factors. Comput Struct Biotechnol J 15:366–371.

47. Tropel D, van der Meer JR. 2004. Bacterial Transcriptional Regulators for Degradation Pathways of Aromatic Compounds. Microbiology and Molecular Biology Reviews 68:474–500.

48. Baugh AC, Defalco JB, Duscent-Maitland C V., Tumen-Velasquez MP, Laniohan NS, Figatner K, Hoover TR, Karls AC, Elliott KT, Neidle EL. 2024. Regulation of tricarboxylate transport and metabolism in *Acinetobacter baylyi* ADP1. Appl Environ Microbiol 90.

49. Uthoff S, Steinbüchel A. 2012. Purification and Characterization of an NAD+-Dependent XylB-like aryl alcohol dehydrogenase identified in *Acinetobacter baylyi* ADP1. Appl Environ Microbiol 78:8743– 8752.

50. Dal S, Trautwein G, Gerischer U. 2005. Transcriptional organization of genes for protocatechuate and quinate degradation from *Acinetobacter* sp. strain ADP1. Appl Environ Microbiol 71:1025–1034.

51. Smith MA, Weaver VB, Young DM, Ornston LN. 2003. Genes for chlorogenate and hydroxycinnamate catabolism (*hca*) are linked to functionally related genes in the *dca-pca-qui-pob-hca* chromosomal cluster of *Acinetobacter* sp. strain ADP1. Appl Environ Microbiol 69:524–532.

52. Parke D, Ornston LN. 2003. Hydroxycinnamate (*hca*) Catabolic Genes from *Acinetobacter* sp. Strain ADP1 Are Repressed by HcaR and Are Induced by Hydroxycinnamoyl-Coenzyme A Thioesters. Appl Environ Microbiol 69:5398.

53. Vries J de, Wackernagel W. 2002. Integration of foreign DNA during natural transformation of *Acinetobacter* sp. by homology-facilitated illegitimate recombination. Proceedings of the National Academy of Sciences 99:2094–2099.

54. Barbe V, Vallenet D, Fonknechten N, Kreimeyer A, Oztas S, Labarre L, Cruveiller S, Robert C, Duprat S, Wincker P, Ornston L, Weissenbach J, Marlière P, Cohen G, Médigue C. 2004. Unique features revealed by the genome sequence of Acinetobacter sp. ADP1, a versatile and naturally transformation competent bacterium. Nucleic Acids Res 32:5766–5779.

55. Metzgar D, Bacher JM, Pezo V, Reader J, Döring V, Schimmel P, Marlière P, de Crécy-Lagard V. 2004. *Acinetobacter* sp. ADP1: An ideal model organism for genetic analysis and genome engineering. Nucleic Acids Res 32:5780–5790.

56. Krahn T, Wibberg D, Maus I, Winkler A, Bontron S, Sczyrba A, Nordmann P, Pühler A, Poirel L, Schlüter A. 2016. Intraspecies Transfer of the Chromosomal *Acinetobacter baumannii* blaNDM-1 Carbapenemase Gene. Antimicrob Agents Chemother 60:3032.

57. Cooper RM, Tsimring L, Hasty J. 2017. Inter-species population dynamics enhance microbial horizontal gene transfer and spread of antibiotic resistance. Elife 6.

58. Heravi KM, Lange J, Watzlawick H, Kalinowski J, Altenbuchner J. 2015. Transcriptional regulation of the vanillate utilization genes (*vanABK* operon) of *Corynebacterium glutamicum* by VanR, a PadR-like repressor. J Bacteriol 197:959–972.

59. Semaming Y, Pannengpetch P, Chattipakorn SC, Chattipakorn N. 2015. Pharmacological Properties of Protocatechuic Acid and Its Potential Roles as Complementary Medicine. Evidence-Based Complementary and Alternative Medicine 2015:1–11.

60. Huang SC, Yen G-C, Chang L-W, Yen W-J, Duh P-D. 2003. Identification of an Antioxidant, Ethyl Protocatechuate, in Peanut Seed Testa. J Agric Food Chem 51:2380–2383.

61. Otsuka Y, Nakamura M, Shigehara K, Sugimura K, Masai E, Ohara S, Katayama Y. 2006. Efficient production of 2-pyrone 4,6-dicarboxylic acid as a novel polymer-based material from protocatechuate by microbial function. Appl Microbiol Biotechnol 71:608–614.

62. Cuff LE, Elliott KT, Seaton SC, Ishaq MK, Laniohan NS, Karls AC, Neidle EL. 2012. Analysis of IS1236-Mediated Gene Amplification Events in *Acinetobacter baylyi* ADP1. J Bacteriol 194:4395.

63. Suárez GA, Renda BA, Dasgupta A, Barrick JE. 2017. Reduced Mutation Rate and Increased Transformability of Transposon-Free *Acinetobacter baylyi* ADP1-ISx. Appl Environ Microbiol 83.

